# Euclidean Distance based Adaptive Sampling Algorithm for Disassociating Transient and Oscillatory Components of Signals

**DOI:** 10.1101/2025.02.23.639754

**Authors:** Safwan Mohammed, Neeraj J. Gandhi, Clara Bourelly, Ahmed Dallal

**Affiliations:** Systems and Biomedical Engineering Department, Faculty of Engineering, Cairo University, Giza, Egypt; Department of Bioengineering, Swanson School of Engineering, University of Pittsburgh, Pittsburgh, PA, USA; Center for Neural Basis of Cognition, University of Pittsburgh, PA; Department of Electrical & Computer Engineering, Swanson School of Engineering, University of Pittsburgh, Pittsburgh, PA, USA

**Keywords:** Adaptive smoothing, neural signals, oscillatory component, signal separation, spectral leakage, transient component, local field potential

## Abstract

Neural signals encode information through oscillatory and transient components. The transient component captures rapid, non-rhythmic changes in response to internal or external events, while the oscillatory component reflects rhythmic patterns critical for processing sensation, action, and cognition. Current spectral and time-domain methods often struggle to distinguish the two components, particularly under sharp transitions, leading to interference and spectral leakage. This study introduces a novel adaptive smoothing algorithm that isolates oscillatory and transient components by dynamically up-sampling signal regions with abrupt changes. The approach leverages Euclidean distance-based thresholds to refine sampling and applies customized smoothing techniques, preserving transient details while minimizing interference. Tested on both synthetic and recorded local field potential data, the algorithm outperformed conventional methods in handling steep signal transitions, as demonstrated by lower mean-square error and improved spectral separation. Our findings highlight the algorithm’s potential to enhance neural signal analysis by more accurately separating components, paving the way for more precise characterization of neural dynamics in research and clinical applications.

## I. Introduction

Continuous neural signals like the electrocorticogram (ECoG), electroencephalogram (EEG), and local field potential (LFP) encode information through oscillatory and transient features. The transient component represents rapid, non-rhythmic activity that typically occurs in response to acute events like stimulus presentation or movement generation. It enables neural circuits to respond adaptively to changes in the environment, signaling the occurrence of new events and contributing to rapid information processing [1, 2, 3]. It often originates from neural processes linked to spikes, especially high frequency bursts, as well as synaptic events leading up to a spike, the low-frequency component of the action potential (“spike bleed-through”), and spike afterhyperpolarization [4]. On the other hand, the oscillatory component arises from network oscillations, rhythmic patterns reflecting structured, synchronized activity across neural populations [3, 5, 6, 7, 8]. Such oscillations are important for transmitting and integrating sensory, motor, and cognitive information.

Spectral methods relying on short-time Fourier transform, multitapers, or wavelet decomposition are the go-to analyses for continuous neural signals [9]. These approaches aim to reveal the frequency content of the signal. However, they often model sharp transitions (i.e., non-oscillatory events) as oscillatory components. This assumption introduces challenges, as transient components create a strong, low-frequency pattern that overshadows the true oscillatory patterns [10, 11], as shown in Figure 1B for LFP data recorded in the monkey superior colliculus. To address this issue, researchers often apply log normalization to reduce the dominance of the low-frequency power over the entire spectrum. Nonetheless, even after normalization, the presence of slowly oscillating components can still dominate over the rhythmic oscillatory events. This interference highlights the limitations of conventional spectral analysis methods, as they fail to fully isolate and accurately represent the underlying oscillatory and transient components.

**Fig. 1.**
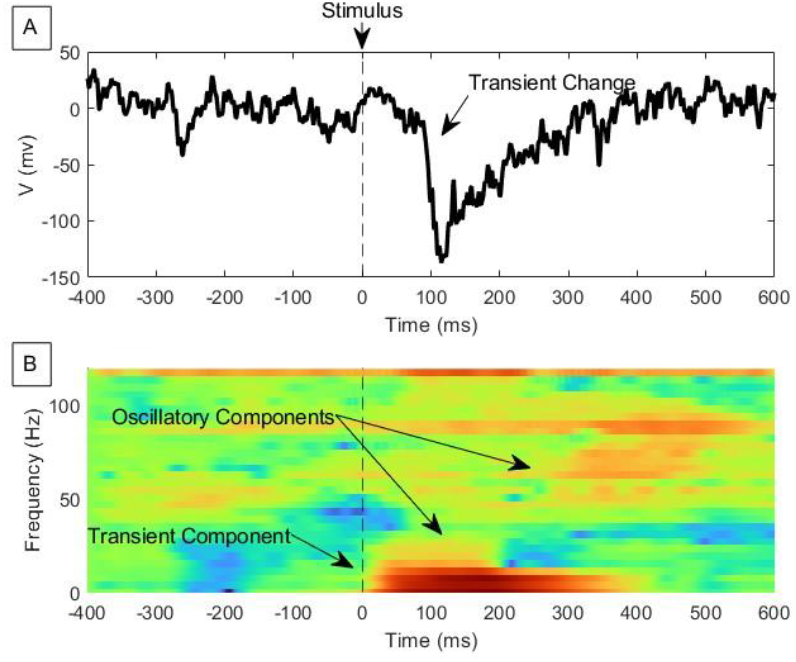
Visual response modulation in LFP from monkey superior colliculus reveals the oscillatory and transient components in the signal. (A) A single-trial LFP signal in the time domain. (B) The corresponding spectrogram using multitapers method.

The Fourier Transform, and other methods built upon it (i.e., Welch and multitapers) [12, 13] while powerful for analyzing the frequency content of signals, struggle with sudden changes or transient modulations. This limitation arises because the transform assumes the signal is periodic and continuous over an infinite duration. When a signal contains abrupt changes, such as sharp edges or transients, the Fourier Transform spreads its frequency representation across multiple components, with higher amplitude affecting low frequencies (1/*f* like shape). These artifacts occur due to the inherent inability of the Fourier Transform to localize changes in both time and frequency domains simultaneously, resulting in a loss of precision when capturing short-lived or non-repetitive features. This limitation makes the Fourier Transform less suitable for analyzing signals with sharp transitions and motivates the need to isolate the transient component from the oscillatory signal [14].

Since the transient change in the signal causes a 1/*f* like component in the frequency domain [15, 16, 17, 18], methods like Fitting Oscillations and One-Over-F (FOOOF) [19] and Irregular-Resampling Auto-Spectral Analysis (IRASA) [20] have been proposed to dissociate the transient component from the oscillatory component (also addressed as aperiodic and periodic components, respectively) using a frequency domain approach. To do so, they estimate the 1/*f* component and subtract it from the Power Spectral Density (PSD) of the original signal. Such methods, however, are very sensitive to the choice of parameters and initial values and can lead to poor fits for the 1/*f* pattern, resulting in false spectral components often with negative power in the oscillatory component [16].

On the other hand, other approaches were proposed to separate the two components in the time domain. Conventional smoothing algorithms like moving average, Savitsky-Golay [21], or local regression can be applied to extract the transient component out of the original signal by smoothing it [17]. However, such methods traditionally use a fixed-size sliding window, which makes them poor at handling rapid or transient changes in amplitude.

In this work, we propose an adaptive method called Euclidean Distance-based Adaptive Sampling (EDAS) for separating the oscillatory and transient components in the time domain. The proposed algorithm employs a customized smoothing technique designed to distinguish oscillatory from non-oscillatory components, effectively isolating features essential for further analysis. The method was tested on both synthetic and neural signals.

## II. METHODS

### A. Dataset

To evaluate the performance of the proposed EDAS algorithm and to compare it against conventional methods, we used both synthetic and real data.

The synthetic signal (*S*_*T*_) was designed to be composed of three components (rhythmic, transient, and noise), and was defined as:

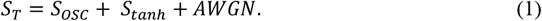

The three components include a rhythmic sinusoidal component (*S*_*0SC*_) featuring three different frequencies, a hyperbolic tangent function (*S*_*tanh*_), and an additive white Gaussian noise (*AWGN*). The hyperbolic tangent function was used to simulate the transient, non-oscillatory change as follows:

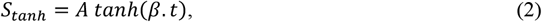

where β is the steepness parameter and it determines the speed of change of the transient component, and A is the amplitude. We assessed and compared the performance of our algorithms for different steepness values.

For the real data, we used LFP recordings from monkey’s superior colliculus during a visually guided eye movement task. All experimental and surgical procedures were approved by the Institutional Animal Care and Use Committee of the University of Pittsburgh and were in accordance with the guidelines of the U.S. Public Health Service policy on the humane care and use of laboratory animals. Experimental details are described in [22]. Briefly, male rhesus monkeys (Macaca mulatta) were seated with their heads restrained in a primate chair and faced a computer monitor. Each animal was trained to direct its visual axis to targets to earn a liquid reward. A multi-contact laminar electrode was concurrently lowered in the superior colliculus to record neural activity. The target was placed at a location relative to the fixation that produced robust modulation in the neural activity. The raw neural signal was sampled at 30 kHz and then low pass filtered (cutoff frequency: 250 Hz) and down-sampled to 1 kHz to yield the LFP. Each single trial trace was then processed through spectral analyses.

### B. Euclidean Distance based Adaptive Sampling (EDAS)

We developed a customized smoothing technique to isolate the oscillatory component of the neural signal from its transient counterpart. Figure 2 represents this algorithm in a block-diagram flow chart form and Figure 3 illustrates the step-by-step application of our algorithm on an LFP recording (Fig. 3A).

**Fig. 2.**
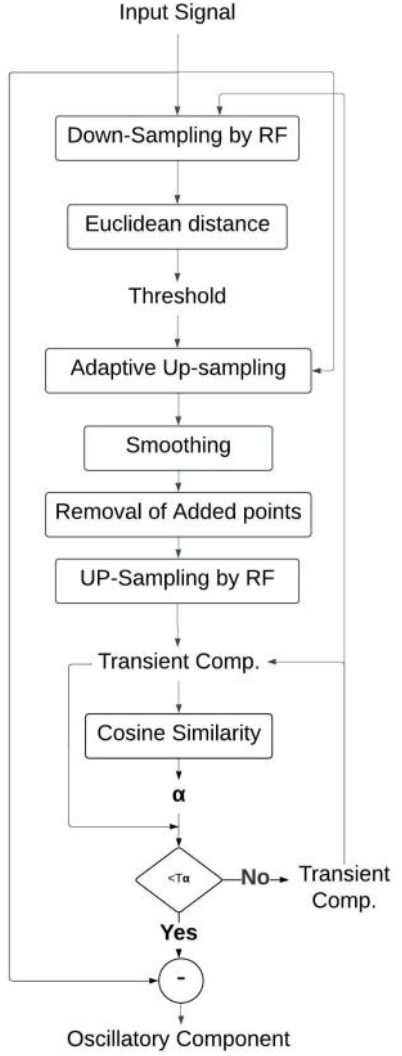
A flow chart representation of EDAS algorithm.

**Fig. 3.**
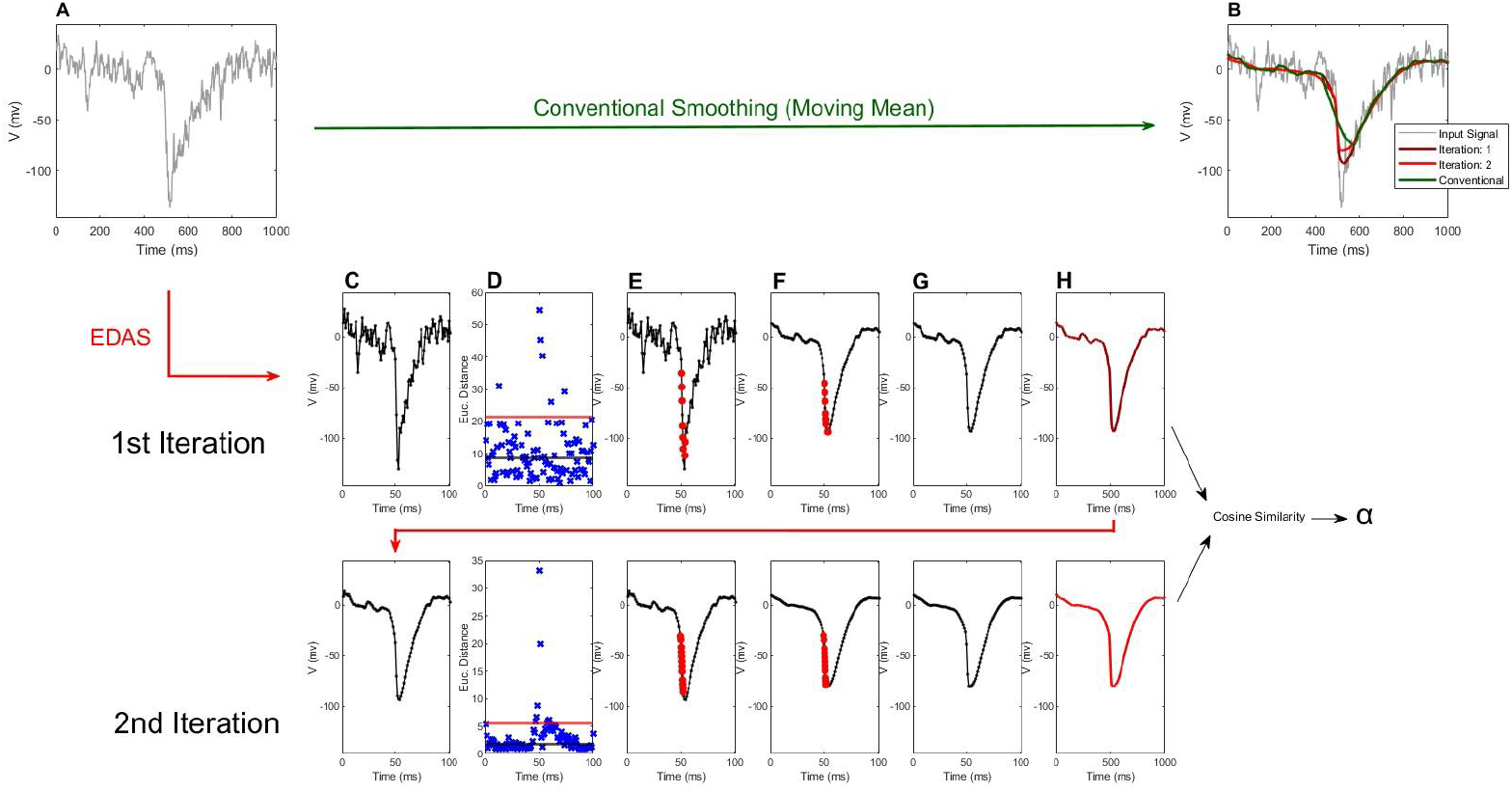
A step-by-step visualization for the EDAS algorithm on an LFP signal, along with the conventional method. (A) Input signal sampled at 1000Hz. (B) Signal smoothed by “conventional” method (moving mean; window size: 0.15 ms). 1st iteration row: (C) Down sampled signal by RF=10 to reduce sampling rate to 100Hz. (D) Euclidean distance between each two consecutive points denoted by blue ‘x’. Black line is the median of distances. Red line denotes threshold *T*_*d*_ that defines which area gets up sampled. (E) Adaptively up sampled signal. Red dots were added as described in text. (F) Smoothed Adaptively up-sampled signal. For this example trial, we computed the moving mean on a 0.15 s window. (G) Smoothed Signal after removing adaptively added samples. New sampling rate is100 Hz, same as in Figure 2C. (H) The signal is then up-sampled to original sampling rate of 1000Hz. The 2nd iteration row performs the same sequence of steps, but the starting point is the output signal sampled at 1000 Hz from the previous iteration. The iteration process continues until the cosine similarity index (*α*), calculated on the outputs of consecutive iterations, surpasses a threshold (*T*_*α*_).

In traditional smoothing algorithms, such as moving average, a window size is defined as a constant span on the abscissa (time) axis, and all points inside this window are averaged. Sharp transitions are characterized by rapid changes in amplitude over a small range of abscissa. When using a fixed window, these transitions may span multiple but limited number of data points within the window, causing the transition’s distinct characteristics to be smoothed out or misrepresented (see Fig. 3B – green trace). To mitigate this, our algorithm adaptively inserts additional data points, i.e., up samples the signal specifically around regions where there is a significant Euclidean distance between consecutive points, relative to the median Euclidean distance between all consecutive point pairs. This ensures that the sharp transition is more finely sampled.

We assume that the time domain signal, *s*(*n*), is decomposed as follows:

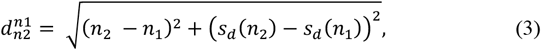

where *H*(*n*) represents the oscillatory component, *NH*(*n*) denotes the transient or non-harmonic component, and *ϵ*(*n*) is zero-mean noise. Our objective is to estimate the transient component and subtract it from the original signal to isolate the oscillatory component for spectral analysis. The EDAS algorithm begins by down-sampling the original signal *s*(*n*) by a Resampling Factor (RF) (Fig. 3C). This step increases the contrast in the distance between consecutive points, especially during sharp transition. Figure 3D shows this distance between consecutive points of the discretized time series. The Euclidean distance between two consecutive samples at time frames *n*_1_ and *n*_2_, is determined as:

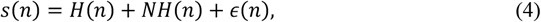

where *s*_*d*_(*n*) is the down-sampled signal in Figure 3C. This distance calculation considers both time and amplitude, allowing the method to detect sudden changes in the signal.

To maintain the characteristics of the transient components, we finely sample the regions experiencing sharp transitions. A threshold, *T*_*d*_, is chosen to ensure up-sampling only those regions that experience sudden changes. We chose:

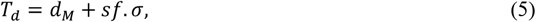

where *d*_*M*_ is the median of calculated distances, and σ is the median absolute deviation, and *sf* (sigma factor) is a tunable parameter controlling *T*_*d*_. The red line in Figure 3D denotes *T*_*d*_ for *sf* = 2. When a large distance is detected (*d*_*i*_ > *T*_*d*_), additional points are inserted using linear interpolation (red dots in Fig. 3E). The number of added points is determined as:

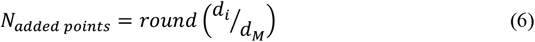

This ensures that regions with abrupt power changes are sufficiently sampled before smoothing. The up-sampled signal, *S*(*n*), is given by:

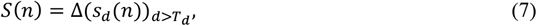

where Δ is the up-sampling operator when the distance exceeds the threshold *T*_*d*_ (Fig. 3E). After adding these points, the signal is smoothed to get the transient component (Fig. 3F). We used moving-mean with a window size of 0.15s for smoothing, but other techniques such as Savitsky-Golay or local regression could also be applied. This computation yields a non-uniformly up-sampled transient component, *NH*_*u*_(*n*), using the moving-mean formula:

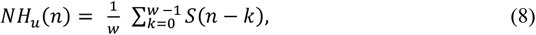

where *w* is the window size. The added samples were then removed to restore the signal to its original uniform sampling (Fig. 3G):

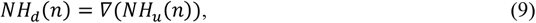

where *∇* is the down-sampling operator. Finally, the estimated transient component, *NH*_*d*_(*n*), is up-sampled back using the same factor RF that was used to down-sample the input signal *s*(*n*) to obtain the estimate of the transient component in *s*(*n*), *i. e*., *NH*(*n*) (Fig. 3H). This transient component is then subtracted from the original signal to estimate the oscillatory component (not shown in Fig. 3).

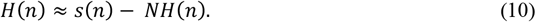

To minimize oscillations in the estimated non-oscillatory component, the algorithm is iteratively repeated, such that the output of iteration (*n −* 1) serves as the input of iteration (*n*), until a stopping condition is met. We defined the stopping criterion through the cosine similarity between the estimated transient components in iterations (*n*) and (*n −* 1). When the cosine similarity is higher than a threshold (which means the transient component did not change much appreciably) we stop the algorithm.

The cosine similarity (*α*) between two signals, x_n_’ and ‘x_n-1_’, was calculated using the following formula:

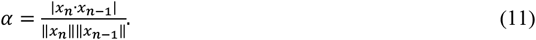

Here, *α* ranges between [-1 1], with 1 representing perfect match, 0 representing no match, and −1 representing perfect inverse match. We set a threshold *T*_*α*_ on *α* (usually 0.99) as a stopping condition. That means when the transient component calculated in this iteration and the transient component from the previous iteration are similar by *T*_*α*_ or more, we stop iterating.

Finally, it is worth noting that changes in some of the algorithm parameters have impacted the convergence of the iterative process. These parameters are listed in Table 1. The values of these parameters were selected manually (see discussion for further information). Given the myriad possibilities, we occasionally set an upper limit on the number of iterations when the convergence rate was slow.

**TABLE 1.** LIST OF TUNABLE OPTIONS AND PARAMETERS.

| Param. | Description |
| --- | --- |
| $W$ | Window size used for smoothing |
| $RF$ | Resampling factor |
| $sf$ | Threshold that defines which area gets up sampled |
| $S\_Method$ | Method used for smoothing |
| $T_\alpha$ | The algorithm stops iterating when $(\alpha > T_\alpha)$ |

## III. Testing and Results

To evaluate the performance of EDAS and analyze its effects and potential complications, several tests were conducted. These tests focused on comparing EDAS against the traditional smoothing methods in both the time and frequency domains.

Objective testing was performed using synthetic data, where the ground truth was known, and the Mean Square Error (MSE) served as the assessment metric. In the time domain, EDAS was applied to the synthetic signal described in Equation (1) alongside multiple smoothing methods, including the moving average, Savitsky-Golay, and local regression. For each method, the MSE was calculated between the method’s estimate of the transient component and the ground truth transient signal, defined as *S*_*tanh*_ in Equation (2), which corresponds to the noiseless transient component. Testing was conducted for various steepness levels, β, of *S*_*tanh*_. Figure 4 illustrates the synthetic signal, along with examples of conventional and EDAS results of the three methods compared to the ground truth (gray solid line). It shows that the conventional methods couldn’t catch the transient change, unlike EDAS. A qualitative assessment demonstrates superior performance by the EDAS algorithm. Figure 5 depicts how different methods react to increasing the steepness of the transient signal. As steepness increases from 0.1 to 1.9, the MSE greatly increases for all three conventional smoothing methods, while EDAS maintains a relatively constant MSE. Paired t-tests were performed between the MSE of EDAS and the MSE of each of the three conventional methods mentioned in Figure 5. All three comparisons showed that the MSE of the EDAS algorithm is significantly smaller than the MSE error from all other traditional methods, (moving average: P<0.001; Savitsky-Golay: P<0.01; local regression: P<0.05). We next used the t-test to assess whether the slope of a regression for each of the traces in Figure 5 is significantly different from zero. All three tests using the conventional method were statistically significant (P<0.001), while none of the three methods used with EDAS had a significant slope (p>0.05). This indicates that EDAS’s MSE is not sensitive to varying steepness levels, thereby supporting the EDAS algorithm’s effectiveness in preserving signal integrity under different conditions. This robustness in performance highlights the importance of the adaptive nature of the algorithm. To study the impact of EDAS in the frequency domain and assess its ability to separate the transient and the oscillatory components, we calculated the transient component of the synthetic data, which contains three sinusoids, using EDAS. Then, the estimated transient component was subtracted from the original signal to obtain the oscillatory component. Fourier transform was calculated for both transient and oscillatory signals and plotted on the same graph. The same procedures were followed for each of the conventional methods. Figure 6 shows the frequency representation of the transient and oscillatory components for the three smoothing methods with (right column) and without (left column) applying EDAS. Results show that conventional smoothing methods caused spectral leakage in the low-frequency band around 5 Hz (less obvious in local regression due to the high-order fitting), which can be misinterpreted as slow oscillations. This effect can be very misleading while working with biological signals, where the oscillatory components usually span a band in the spectral domain, rather than just an impulse-shaped spike. However, EDAS showed a nearly perfect isolation between the transient and the oscillatory components in the spectral domain.

**Fig. 4.**
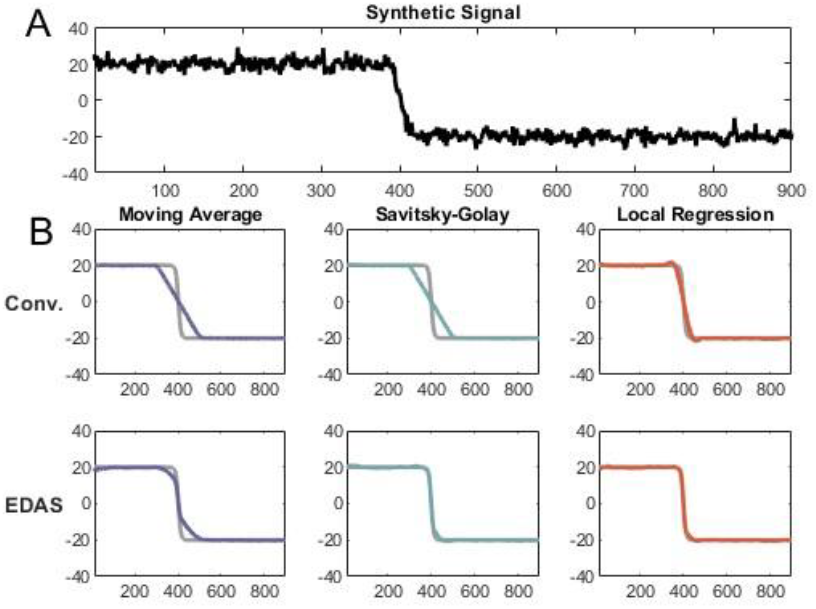
(A) Synthetic data used for testing, composed of a hyperbolic tangent, three sinusoids and zero-mean noise. (B) Smoothed signal with conventional methods (top row) and EDAS (bottom row) for moving average, Savitsky-Golay, and local regression, plotted with noiseless transient component *S*_*tanh*_ in equation 2.

**Fig. 5.**
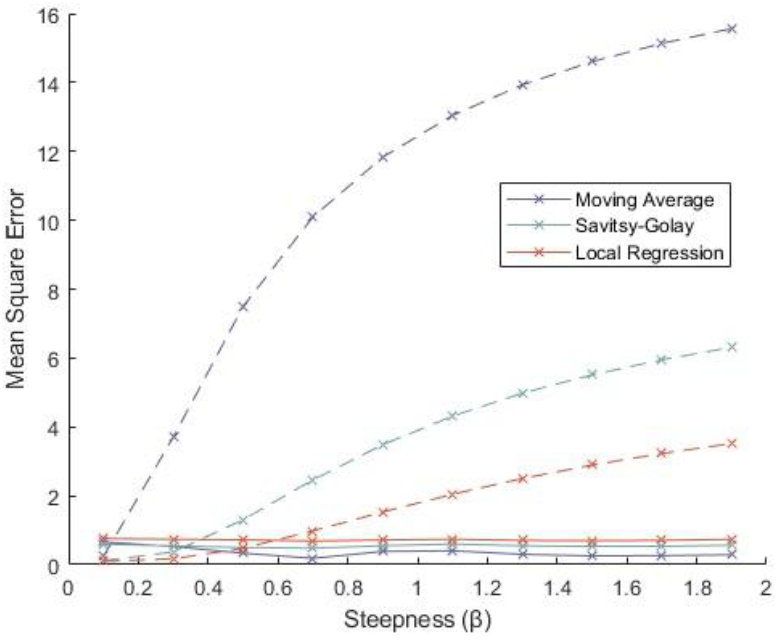
MSE vs. the Steepness of the transient signal computed for conventional against proposed method for moving average, Savitsky-Golay, and local regression. Continuous line represents the proposed method, dashed line represents conventional method.

**Fig. 6.**
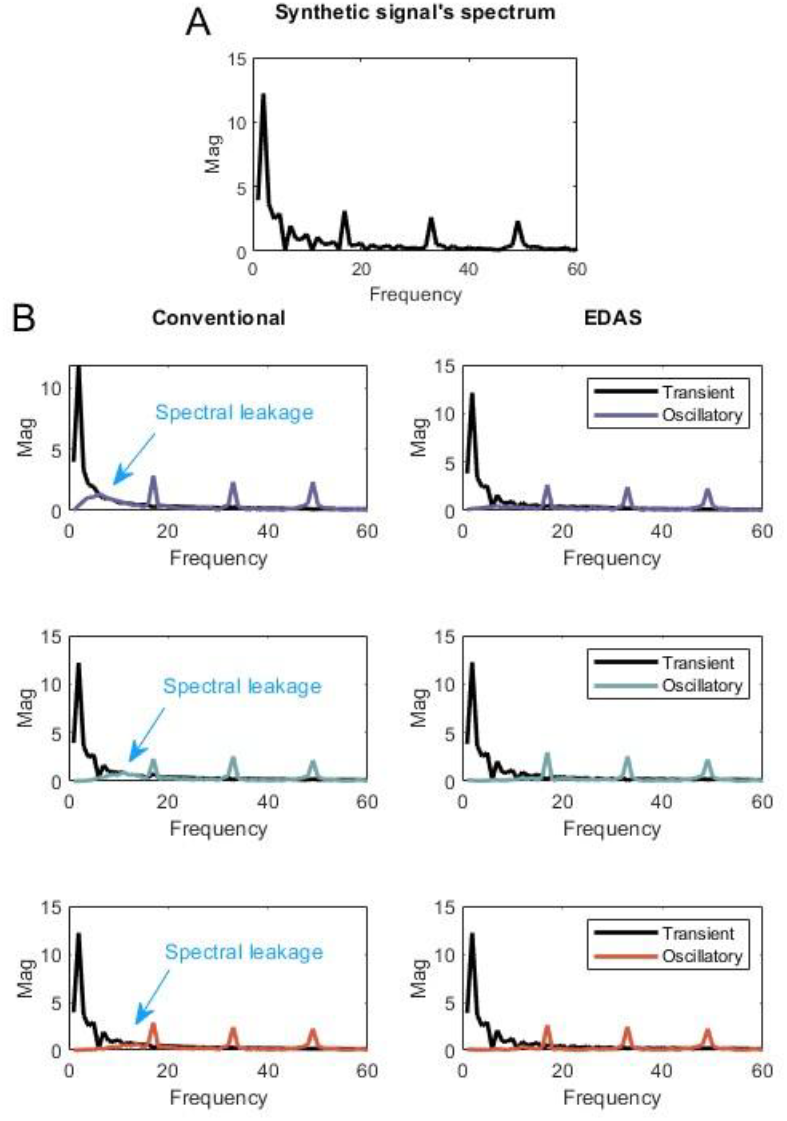
(A) Spectrum of synthetic signal *S*_*T*_ calculated by Fourier transform (FFT) (B) Spectral shape calculated by FFT for Transient and Oscillatory component for conventional (left column) and proposed (right column) methods. First raw: moving average. Second raw: Savitsky-Golay. Third raw: Local regression.

To explore the effect of the proposed algorithm on real data, LFP signals were recorded from the superior colliculus of a monkey producing visually guided saccadic eye movements. Figure 7 shows the raw signal (A) from a single trial and its corresponding spectrogram (B), as well as the transient (C) and the oscillatory components (E) isolated by EDAS. By comparing Figures 7A and 7C, one can notice that the algorithm managed to adapt to the sudden change (300-500 ms) without overfitting to the oscillations in the other areas. The isolation can also be observed in the spectrogram (Fig. 7D) as the transient component exhibits a high-power low frequency footprint, which can be misleading while interpreting biological signals, if not isolated.

**Fig. 7.**
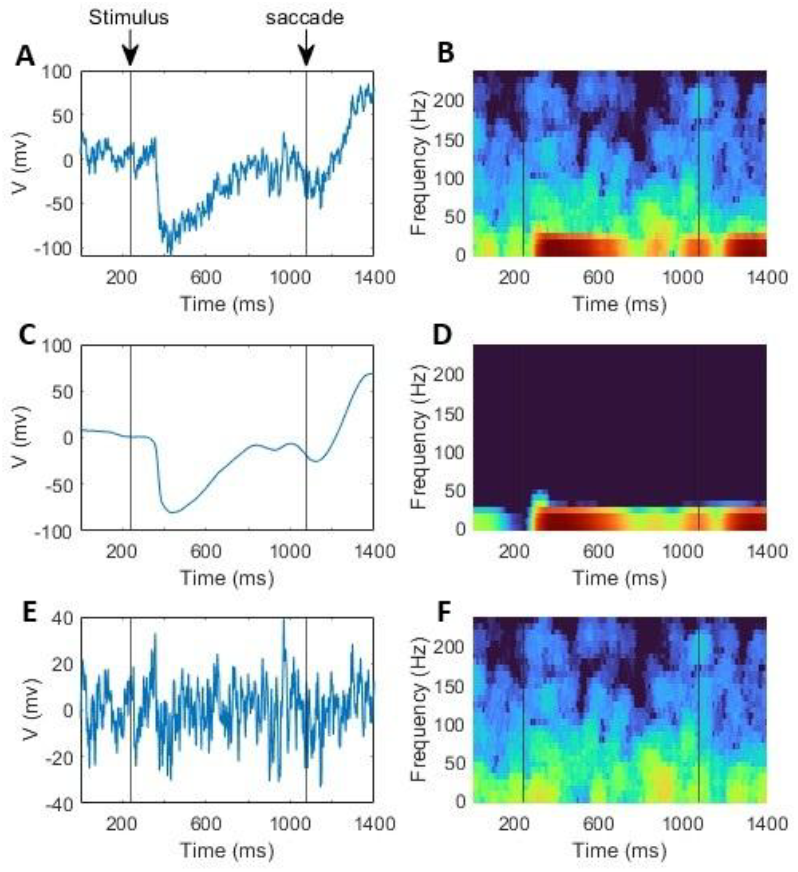
Separating LFP signal into transient and oscillatory components, and the effect on spectrogram. The three spectrograms were subjected to log normalization, and use the same color scale.

We executed the algorithm on 10 experiments, comprising a total of 38,192 waveforms, with *T*_*α*_ = 0.99. The number of iterations for each trial was noted (mean = 2.7, std = 0.5), with ~70% converging in fewer than 4 iterations.

## IV. Discussion

In this study, we developed an Euclidean Distance-based Adaptive Sampling (EDAS) algorithm to address the challenge of separating the transient and oscillatory components in time series signals, and we extended its application to neural data [22]. Traditional methods, including spectral and fixed-window smoothing approaches, struggle with accurately distinguishing these components, particularly during sharp transition periods, resulting in spectral leakage and misrepresentation of neural dynamics. Our algorithm mitigates these concerns by dynamically up-sampling signal regions with significant Euclidean distance changes and applying adaptive sampling techniques to preserve transient details while isolating oscillatory features.

### A. Adaptive Smoothing Techniques in Signal Processing

Efficient smoothing of signals with transient changes has been a focus in recent research, with notable contributions from the Multiscale Polynomial Filter (MPF) [23] and Extreme Envelope Average (EEA) [24] algorithms. As an extension of the Savitsky-Golay filter, the MPF adapts the smoothing window based on local signal complexity and noise. Unlike fixed-window SG filters, the MPF adjusts its window size dynamically to balance noise reduction and signal fidelity. However, its reliance on polynomial fitting can introduce bias and excessive smoothing in regions with abrupt transitions. It is, however, susceptible to bias in high-curvature regions, leading to loss of sharp details. The EEA reduces noise by iteratively averaging the signal’s maximum and minimum envelopes. It excels at preserving low-frequency components but struggles with sharp transitions and high-frequency events, often requiring fine-tuning for noisy signals. It struggles with precise transient resolution and convergence in high-noise situations. On the other hand, the proposed EDAS algorithm offers several key advantages, making it highly efficient for signal processing and analysis. It excels in handling sharp transition by identifying and up-sampling regions with significant changes, ensuring that transient features are preserved. Additionally, its high temporal resolution allows it to capture fast, transient events without dilution, maintaining the integrity of rapid signal variations. These features make EDAS a powerful tool for applications requiring precision and adaptability.

### B. Literature Insights on Neural Signal Components

Spectral analysis of LFP signals is a fundamental method in systems neuroscience. It is widely used to assess frequency content relative to baseline periods. Conventional techniques, such as multitapers spectrograms, have been essential for identifying dominant rhythms within neural processes, with beta (13–30 Hz) and gamma (30–100 Hz) frequency bands emerging as crucial biomarkers associated with a concomitant effect on sensation, action, and cognition as well as impairments in neuropathological conditions [25, 26]. Beta frequency oscillations play a central role in the coordination and execution of movements by modulating neural communication across motor circuits [27, 28]. Beta power is persistently elevated in default states, such as baseline and periods of motor preparation and postural maintenance, but it decreases during disruptions or transitions such as initiating a new movement or shifting attention [29]. Following movement completion, beta oscillations can rebound, reflecting a return to a stable, regulated state. On the other hand, gamma frequency modulation is critical for cognitive function. Gamma power is elevated when stimuli are behaviorally relevant or attended, suggesting that they help filter and prioritize relevant input while suppressing noise [25]. It is also crucial in top-down processes, as attention and working memory demands can modulate gamma amplitude and coherence, reflecting a mechanism by which higher-order regions exert influence on local sensory and motor circuits [30].

Transforming a time series signal into the frequency domain assumes that the signal is stationary. The LFP indeed has an underlying component whose statistical properties obey this requirement. However, the LFP also integrates transient features that can be linked to specific and brief features like stimulus presentation or movement generation. In other words, the time domain signal is composed of both oscillatory and transient components and dissociating them has become a focus of recent studies. One approach has been to estimate and remove the transient component in the frequency domain, through methods like FOOOF and IRASA [19, 20]. Another framework, like the proposed EDAS method, removes the nonharmonic component in the time domain. The proposed approach offers a robust framework to disentangle transient and oscillatory components by enhancing temporal resolution and preserving transient dynamics. Its adaptive sampling approach improves time-domain precision while minimizing distortion, making it a powerful alternative to conventional spectral methods. By isolating neural components with greater accuracy, EDAS responds to the demand for innovative tools in neuroscience, advancing the analysis of complex neural signals. In our experiments, the EDAS method qualitatively captured the features of multitapers spectrograms in frequencies above the beta band, while substantially removing lower frequency components (Fig. 7).

Transient changes associated with stimulus onset and movement generation are pronounced and relatively easy to detect in LFP signals recorded from a region involved in sensation and action. In Figure 1, for example, the abrupt drop in voltage around 100 ms reflects the registration of a stimulus present at time 0, with the delay attributed to transduction time from the eye to the superior colliculus. However, transient modulations, usually of smaller amplitude and last <150 ms, can occur at different times and rates and at varying intensities on individual trials [31, 32, 33, 34, 35]. For example, Figure 1A shows a deflection at −250 ms and potentially another at +350 ms. It may not be straightforward to link the transient to a behavior. Thus, caution must be employed when determining whether such transient changes should be part of the nonharmonic or oscillatory signal. One or more tuning parameters (see Table 1) may need to be modified to achieve the desired result.

### C. EDAS in neural applications

Neural signals are characterized by an aperiodic component, often called 1/*f* or 1/*f*^*β*^, which typically manifests as a 1/*f* shaped PSD. However, this shape can vary, showing features like a “knee” in the low-frequency range or a spectral plateau at higher frequencies [13]. The exact origin of the 1/*f* component remains uncertain. Methods such as FOOOF attempt to fit a predefined 1/*f*^*β*^ function, with modifications to account for features like the knee. However, these methods often struggle to fit the entire PSD effectively [16]. As a result, researchers tend to focus on narrow frequency bands and overlook the knee or plateau regions.

In our study, the PSD of certain neural signals exhibited a distinct knee in the low-frequency range. This complexity caused the FOOOF fitting process to fail when using standard initial conditions, requiring extensive manual adjustments (analysis not shown). This limitation made the approach impractical for calculating spectrograms, which involve processing numerous time windows for each signal.

We propose that the 1/*f*^*β*^ component may originate from multiple sources, one of which is the Gibbs phenomenon. This phenomenon arises when a signal undergoes a transient change, producing a sinc-shaped frequency response. These transient changes often correspond to Event-Related Potentials (ERPs). One common method for addressing this involves averaging multiple trials to extract the ERP and then subtracting it from each trial. However, this approach has significant limitations. It assumes that oscillations will cancel out across trials, which may not always happen. Oscillations could synchronize across trials, leading to the unintentional removal of critical oscillatory activity from each trial. To address these issues, we recommend using the EDAS method for tasks involving event-driven signals, such as those in stimulus-response or motor task experiments.

### D. Future Work

While the EDAS algorithm has demonstrated its efficacy in separating transient and oscillatory components, there remains significant potential for further refinement and enhancement. The current implementation is limited by the manual selection of tunable parameters and using the cosine similarity as a stopping condition. A promising direction for future work involves formulating the separation process as an optimization problem. By defining a loss function that quantifies the quality of separation—potentially incorporating measures such as spectral leakage, transient preservation, and oscillatory integrity—we can leverage optimization algorithms to automate the tuning of parameters.

This approach would offer several advantages:

1. *Automated Parameter Selection:* Current methods require manual tuning of parameters such as the threshold for up sampling and smoothing window sizes. An optimization framework would eliminate this reliance, allowing for dynamic and data-driven adjustments tailored to each signal.

2. *Increased Robustness and Confidence:* By systematically minimizing the loss function, the optimization process can ensure consistency and reliability in the separation outcomes, reducing the subjectivity introduced by manual parameter selection.

3. *Broader Applicability:* An optimized EDAS algorithm could adapt to a wider variety of neural signal datasets, including those with varying noise levels, transient characteristics, and oscillatory patterns, thereby enhancing its utility across diverse applications in neuroscience research.

4. Incorporating optimization techniques into the EDAS framework would not only improve its performance but also contribute to advancing methodologies for neural signal analysis, enabling more precise and automated characterization of complex neural dynamics.

In summary, the proposed method offers a targeted approach that maintains oscillatory and transient components separately, ensuring high fidelity in neural signal representation. By combining adaptive up sampling with tailored smoothing, it addresses the shortcomings of traditional methods and provides a more robust solution for complex, rapidly varying signals.

## V. Conclusion

In this study, we introduced a novel, Euclidean Distance-based Adaptive Sampling (EDAS) algorithm to separate transient and oscillatory components in signals. Traditional spectral and time-domain methods often struggle with accurately distinguishing these components, particularly during sharp transitions. By dynamically up-sampling signal regions with abrupt changes and applying adaptive smoothing techniques, EDAS extracts transient details while preserving oscillatory features. We showed that EDAS outperforms conventional smoothing methods on both synthetic and real neural data. Future work should focus on automating the parameter selection process and incorporating optimization techniques to further refine the algorithm’s performance. By formulating the separation process as an optimization problem, we can reduce the reliance on manual tuning and improve the algorithm’s adaptability to diverse neural datasets. This would not only enhance the robustness of EDAS but also broaden its applicability across different research and clinical contexts.

## Appendix

A MATLAB implementation of EDAS can be found in this [GitHub repository](https://github.com/Safwanmahmoud/EDAS), along with an example of how to use it.

## Acknowledgment

Research was supported by NIH grant R01 EY024831 (NJG).

## References

[1] C. Wiest, F. Torrecillos, A. Pogosyan, M. Bange, M. Muthuraman, S. Groppa, N. Hulse, H. Hasegawa, K. Ashkan, F. Baig, F. Morgante, E. A. Pereira, N. Mallet, P. J. Magill, P. Brown, A. Sharott, and H. Tan, “The aperiodic exponent of subthalamic field potentials reflects excitation/inhibition balance in Parkinsonism,” Elife, vol. 12, Feb 22, 2023.

[2] N. Adelhofer, T. Paulus, M. Muckschel, T. Baumer, A. Bluschke, A. Takacs, E. Toth-Faber, Z. Tarnok, V. Roessner, A. Weissbach, A. Munchau, and C. Beste, “Increased scale-free and aperiodic neural activity during sensorimotor integration-a novel facet in Tourette syndrome,” Brain Commun, vol. 3, no. 4, pp. fcab250, 2021.

[3] K. Thuwal, A. Banerjee, and D. Roy, “Aperiodic and Periodic Components of Ongoing Oscillatory Brain Dynamics Link Distinct Functional Aspects of Cognition across Adult Lifespan,” eNeuro, vol. 8, no. 5, Sep-Oct, 2021.

[4] S. Ray, “Challenges in the quantification and interpretation of spike-LFP relationships,” Curr Opin Neurobiol, vol. 31, pp. 111–8, Apr, 2015.

[5] G. Buzsaki, Rhythms of the Brain: Oxford university press, 2006.

[6] N. Cortes, B. Oliveira, and C. Casanova, “AB003. Pulvinar mediates the transmission of gamma and alpha oscillatory bandsbetween areas 17 and 21a,” Annals of Eye Science, vol. 4, pp. AB003–AB003, 2019.

[7] S. Cole, and B. Voytek, “Cycle-by-cycle analysis of neural oscillations,” J Neurophysiol, vol. 122, no. 2, pp. 849–861, Aug 1, 2019.

[8] B. Zhang, H. Tian, Y. Yu, X. Zhen, L. Zhang, Y. Yuan, and L. Wang, “A localized pallidal physiomarker in Meige syndrome,” Front Neurol, vol. 14, pp. 1286634, 2023.

[9] B. Pesaran, “Spectral analysis for neural signals,” Short Course III, Society for Neuroscience, vol. 1, 2008.

[10] D. Gottlieb, and C.-W. Shu, “On the Gibbs phenomenon and its resolution,” SIAM review, vol. 39, no. 4, pp. 644–668, 1997.

[11] T. Donoghue, J. Dominguez, and B. Voytek, “Electrophysiological Frequency Band Ratio Measures Conflate Periodic and Aperiodic Neural Activity,” eNeuro, vol. 7, no. 6, Nov-Dec, 2020.

[12] P. Welch, “The use of fast Fourier transform for the estimation of power spectra: A method based on time averaging over short, modified periodograms,” IEEE Transactions on audio and electroacoustics, vol. 15, no. 2, pp. 70–73, 2003.

[13] D. J. Thomson, “Spectrum estimation and harmonic analysis,” Proceedings of the IEEE, vol. 70, no. 9, pp. 1055–1096, 2005.

[14] B. D. Shizgal, and J.-H. Jung, “Towards the resolution of the Gibbs phenomena,” Journal of Computational and Applied Mathematics, vol. 161, no. 1, pp. 41–65, 2003.

[15] K. J. Miller, L. B. Sorensen, J. G. Ojemann, and M. den Nijs, “Power-law scaling in the brain surface electric potential,” PLoS Comput Biol, vol. 5, no. 12, pp. e1000609, Dec, 2009.

[16] M. Gerster, G. Waterstraat, V. Litvak, K. Lehnertz, A. Schnitzler, E. Florin, G. Curio, and V. Nikulin, “Separating Neural Oscillations from Aperiodic 1/f Activity: Challenges and Recommendations,” Neuroinformatics, vol. 20, no. 4, pp. 991–1012, Oct, 2022.

[17] D. Vidaurre, R. M. Cichy, and M. W. Woolrich, “Dissociable Components of Information Encoding in Human Perception,” Cereb Cortex, vol. 31, no. 12, pp. 5664–5675, Oct 22, 2021.

[18] R. M. Borah, A. Pathak, and A. Banerjee, “Emotion arousal but not valence is strongly represented in aperiodic EEG activity stemming from thalamocortical interactions,” bioRxiv, pp. 2024.03.11.584477, 2024.

[19] T. Donoghue, M. Haller, E. J. Peterson, P. Varma, P. Sebastian, R. Gao, T. Noto, A. H. Lara, J. D. Wallis, R. T. Knight, A. Shestyuk, and B. Voytek, “Parameterizing neural power spectra into periodic and aperiodic components,” Nat Neurosci, vol. 23, no. 12, pp. 1655–1665, Dec, 2020.

[20] H. Wen, and Z. Liu, “Separating Fractal and Oscillatory Components in the Power Spectrum of Neurophysiological Signal,” Brain Topogr, vol. 29, no. 1, pp. 13–26, Jan, 2016.

[21] A. Savitzky, and M. J. Golay, “Smoothing and differentiation of data by simplified least squares procedures,” Analytical chemistry, vol. 36, no. 8, pp. 1627–1639, 1964.

[22] C. Bourrelly, C. Massot, and N. J. Gandhi, “Rapid Input-Output Transformation between Local Field Potential and Spiking Activity during Sensation but not Action in the Superior Colliculus,” J Neurosci, vol. 43, no. 22, pp. 4047–4061, May 31, 2023.

[23] M. Browne, N. Mayer, and T. R. Cutmore, “A multiscale polynomial filter for adaptive smoothing,” Digital Signal Processing, vol. 17, no. 1, pp. 69–75, 2007.

[24] F. B. Batista, “One-dimensional smoothing using extreme envelope average,” Mechanical systems and signal processing, vol. 28, pp. 432–442, 2012.

[25] S. Ray, and J. H. Maunsell, “Different origins of gamma rhythm and high-gamma activity in macaque visual cortex,” PLoS Biol, vol. 9, no. 4, pp. e1000610, Apr, 2011.

[26] A. K. Engel, P. Fries, and W. Singer, “Dynamic predictions: oscillations and synchrony in top-down processing,” Nat Rev Neurosci, vol. 2, no. 10, pp. 704–16, Oct, 2001.

[27] B. E. Kilavik, M. Zaepffel, A. Brovelli, W. A. MacKay, and A. Riehle, “The ups and downs of beta oscillations in sensorimotor cortex,” Exp Neurol, vol. 245, pp. 15–26, Jul, 2013.

[28] P. Brown, “Abnormal oscillatory synchronisation in the motor system leads to impaired movement,” Curr Opin Neurobiol, vol. 17, no. 6, pp. 656–64, Dec, 2007.

[29] A. K. Engel, and P. Fries, “Beta-band oscillations--signalling the status quo?,” Curr Opin Neurobiol, vol. 20, no. 2, pp. 156–65, Apr, 2010.

[30] P. Fries, “A mechanism for cognitive dynamics: neuronal communication through neuronal coherence,” Trends Cogn Sci, vol. 9, no. 10, pp. 474–80, Oct, 2005.

[31] H. Shin, R. Law, S. Tsutsui, C. I. Moore, and S. R. Jones, “The rate of transient beta frequency events predicts behavior across tasks and species,” Elife, vol. 6, Nov 6, 2017.

[32] G. Tinkhauser, A. Pogosyan, S. Little, M. Beudel, D. M. Herz, H. Tan, and P. Brown, “The modulatory effect of adaptive deep brain stimulation on beta bursts in Parkinson’s disease,” Brain, vol. 140, no. 4, pp. 1053–1067, Apr 1, 2017.

[33] M. Lundqvist, J. Rose, P. Herman, S. L. Brincat, T. J. Buschman, and E. K. Miller, “Gamma and beta bursts underlie working memory,” Neuron, vol. 90, no. 1, pp. 152–164, Apr 6, 2016.

[34] S. R. Jones, “When brain rhythms aren’t ‘rhythmic’: implication for their mechanisms and meaning,” Curr Opin Neurobiol, vol. 40, pp. 72–80, Oct, 2016.

[35] J. Feingold, D. J. Gibson, B. DePasquale, and A. M. Graybiel, “Bursts of beta oscillation differentiate postperformance activity in the striatum and motor cortex of monkeys performing movement tasks,” Proc Natl Acad Sci U S A, vol. 112, no. 44, pp. 13687–92, Nov 3, 2015.

